# Structural characterization of ribosome recycling and fusidic acid inhibition in *Staphylococcus aureus*

**DOI:** 10.64898/2026.04.27.721003

**Authors:** Adrián González-López, Maria Selmer

**Affiliations:** Department of Cell and Molecular Biology, Uppsala University, Uppsala, 75124, Sweden; Uppsala Antibiotic Center, Uppsala University, Uppsala, 75124, Sweden

## Abstract

During bacterial ribosome recycling, 70S ribosomes are split into subunits by ribosome recycling factor (RRF) and elongation factor G (EF-G). The antibiotic fusidic acid (FA) inhibits elongation and ribosome recycling by locking EF-G to the ribosome. Yet, no functional ribosome recycling FA complex has been successfully captured. Here we used single-particle cryo-electron microscopy to resolve multiple FA-stalled intermediates of Staphylococcus aureus ribosomes, including a previously unobserved complex containing both RRF and EF-G. Our structures reveal how RRF and EF-G jointly disrupt inter-subunit bridges, promote back-rotation of the small subunit, and destabilize the tRNA to facilitate ribosome splitting. We further show that FA predominantly inhibits recycling by trapping EF-G on the post-termination complex in the absence of RRF, preventing formation of the active RRF•EF-G complex. These insights advance understanding of the molecular mechanism of bacterial ribosome recycling and the mode of action of FA as an antibiotic.

**GRAPHICAL ABSTRACT:** 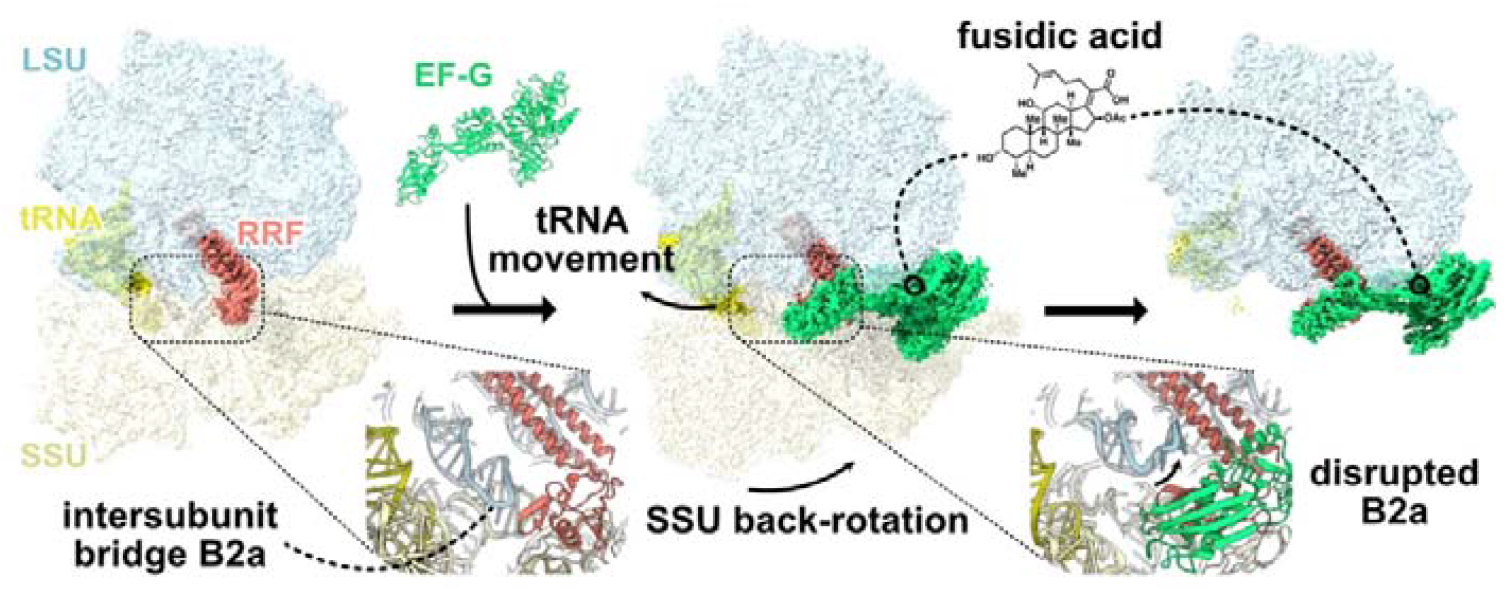

## INTRODUCTION

Ribosome recycling is the final step of bacterial protein synthesis, following translation initiation, elongation and termination. After termination, the ribosome remains as a 70S ribosome with a deacylated P-site tRNA and an empty A site, forming what is known as the post-termination complex (postTC). Ribosome recycling factor (RRF) and elongation factor G (EF-G) act together to dissociate this 70S ribosome into its subunits in a GTP-dependent manner (1), allowing the subunits to participate in new rounds of initiation (2). This splitting process requires the disruption of several inter-subunit bridges, where bridge B2a, formed between helix 44 (h44) of 16S RNA and H69 of 23S RNA near the decoding center, has been suggested as central to the mechanism (3, 4). While the structural mechanism of EF-G-mediated translocation is well understood (5–7), several aspects of ribosome recycling remain unresolved, as recently reviewed (8).

RRF is an L-shaped protein composed of a three-helix bundle (domain I) and an a/b-domain (domain II) (9, 10). The postTC adopts a rotated state (11), to which RRF binds with domain I spanning the A and P-sites (4, 12), while domain II in two possible orientations (13). This binding is only compatible with a deacylated P-site tRNA (14), while in absence of tRNA, RRF can also bind to classical-state ribosomes (15, 16). In some structures, RRF alone induces rearrangements in H69 (15, 17), whereas others detect no such changes (16). The RRF binding site overlaps with the post-translocational position of EF-G, consistent with the requirement that RRF must bind first during recycling (18). Upon EF-G binding and GTP hydrolysis, domain II of RRF rotates, a movement proposed to drive ribosome splitting (19). Low-resolution time-resolved cryo-EM studies suggested that, in the presence of EF-G, domain II of RRF moves towards inter-subunit bridge B2a (13), but the density was of insufficient quality to unambiguously place domain II of RRF. Therefore, there is still no direct structural observation of how RRF together with EF-G breaks essential inter-subunit bridges. It also remains unclear how these structural changes affect tRNA and mRNA to allow their release (8, 20).

The antibiotic fusidic acid (FA) has been successfully used to visualize EF-G on the ribosome (5, 21, 22). It binds to EF-G and prevents the conformational changes required for EF-G release after GTP hydrolysis and translocation (23, 24). FA also inhibits recycling (25, 26), where it can trap EF-G on the postTC in presence or absence of RRF (26). No structure has yet captured a functionally engaged EF-G locked by FA together with RRF on the ribosome.

Here, we use cryo-EM in the presence of FA to resolve multiple ribosome recycling complexes from the clinically relevant bacterium *Staphylococcus aureus* (27). The structures clarify the fundamental mechanism of bacterial ribosome recycling and reveal how FA perturbs this essential process. We visualize a previously unseen intermediate containing both EF-G and RRF bound to the ribosome, providing direct structural evidence for how their coordinated actions destabilize inter-subunit bridges, induce back-rotation of the ribosome and perturb tRNA-30S interactions.

## MATERIAL AND METHODS

### Cloning, overexpression and purification of S. *aureus* EF-G and RRF

*S. aureus* EF-G was purified as published (22) through immobilized metal chromatography and size exclusion chromatography. The final protein sample was stored in 50 mM Tris-HCl pH 7.5, 300 mM NaCl, 5 mM b-mercaptoethanol at -70 °C. This construct includes a 6xHis tag with a TEV protease cleavage site and a linker to the EF-G sequence. After TEV cleavage, only Ser-Leu remains at the N-terminus of EF-G.

The *S. aureus RRF* gene was ordered by gene synthesis from GenScript (Piscataway, New Jersey, United States) in pET28a(+)-TEV. This construct encodes a 6xHis tag with a TEV protease cleavage site and a linker to the RRF sequence. After TEV cleavage, only Gly-His remains at the N-terminus. The plasmid was transformed into *Escherichia coli* BL21 (DE3) and an overnight culture was inoculated 1:100 into a 2.8 l baffled flask with 800 mL of LB with 50 µg/mL of kanamycin. Protein expression was induced at an OD600 of 0.5-0.6 with 1 mM isopropyl-b-thiogalactopyranoside and the culture was incubated for 3 h at 37 °C. The cells were harvested at 8 000 x g for 15 min in a JLA 9.1000 rotor (Beckman Coulter, Brea, California, USA), washed with 150 mM NaCl, and stored at -20 °C until use. Next, the cells were resuspended in lysis buffer (20 mM Tris-HCl pH 7.5, 500 mM NaCl, 1 mM MgSO_4_ 10 mM imidazole, 10 % glycerol) with 0.1 % (v/v) Triton X-100, one EDTA-free mini-Complete protease inhibitor tablet (Roche, Basel, Switzerland) and DNase I. The resuspended cells were lysed in a flow cell disruptor (Constant Systems Ltd., Daventry, United Kingdom) and the lysate was centrifuged at 40 000 x g for 30 min in a JA-25.50 rotor (Beckman Coulter).

The supernatant was filtered by 0.45 µm with a polyethersulfone syringe filter (Sarstedt AG & Co, Nümbrecht, Germany) and incubated with 1 mL of Protino® Ni-NTA Agarose (MACHEREY-NAGEL, Düren, Germany) pre-equilibrated with lysis buffer. The column was washed with lysis buffer until baseline and the protein was eluted with elution buffer (20 mM Tris pH 7.5, 300 mM NaCl, 1 mM MgSO_4_, 500 mM imidazole). The eluate was exchanged into GF buffer (20 mM Tris-HCl pH 7.5, 300 mM NaCl, 1 mM MgSO_4_) through a dialysis membrane 3.5 kDa cut-off Spectra/Por® (Spectrum Laboratory Products Inc., New Brunswick, New Jersey, USA) in presence of 1:50 molar ratio of TEV protease for 16-18 h at 4 °C. This was followed by reverse immobilized affinity chromatography on 250 µL of Ni-NTA Agarose. RRF was further purified by gel filtration on a Hiload 16/60 Superdex-75 column (GE Healthcare, Uppsala, Sweden) pre-equilibrated with GF buffer. The peak fractions were subjected to additional reverse IMAC with 1 ml of Ni Sepharose Fast Flow to remove any uncleaved protein. The final sample was concentrated to 18 mg/mL with a 10-kDa cutoff Vivaspin Turbo 15 (Sartorius AG, Göttingen, Germany), frozen in liquid nitrogen, and stored at -70 °C.

### Ribosome and tRNA purification

S. aureus NCTC 8325-4 ribosomes were purified as published (22). In short, cells were harvested at OD600 of 1.0, lysed with lysostaphin, and pelleted. The supernatant was centrifuged through a 1.1 M sucrose cushion twice. The resulting crude ribosome pellet was resuspended and 70S ribosomes were isolated through rate-zonal centrifugation in a preformed 10-40 % sucrose gradient. The 70S ribosomes were pooled and pelleted. The final 70S sample was stored in HEPES polymix buffer (5□mM HEPES pH 7.5, 5□mM NH_4_Cl, 5□mM Mg(OAc)_2_, 100□mM KCl, 0.5□mM CaCl_2_, 8□mM putrescine, 1□mM spermidine, and 1□mM dithioerythritol) at −70 °C. mRNA Z4AUGAGC (5’-GGCAAGGAGGUAAAAAUG**AGC**AAA -3’) was produced by chemical synthesis (GenScript) and *E. coli* tRNA^Ser^ was overexpressed and purified as published (28).

### Post-termination complex preparation

The sample was prepared by mixing 3 µM (final concentrations) 70S *S. aureus* ribosomes in HEPES polymix buffer (with 20 mM HEPES pH 7.5 and 5 mM BME as reducing agent) with 6 µM mRNA Z4AUGAGC and incubated for 10 min at 37 °C. Then, 6 µM *E. coli* tRNA^Ser^ was added and the mix was incubated for 10 min at 37°C. Next, 40 µM *S. aureus* RRF was added, incubated for 10 min at 37 °C and followed by adding 1 mM GTP. The post-termination complex was kept on ice until vitrification.

### Cryo-EM grid preparation

2.4 µL of 40 µM EF-G with 2 mM GTP and 500 µM FA was mixed with 6 µL of Post-termination complex on ice, incubated 45 s and 3 µl of were deposited on a glow-discharged (15 s at 20 mA) QuantiFoil 200-mesh R 2/2 copper grid with 3 nm continuous carbon, incubated 10 s, blotted for 3 s and plunge-frozen in a Vitrobot Mark IV (Thermo Fisher Scientific, Waltham, MA, USA) at 4 °C and 95 % humidity.

### Cryo-EM data collection

All the grids were screened on a Glacios TEM operated at 200 kV equipped with a Falcon-4i direct electron detector with Selectris energy filter (Thermo Fisher Scientific). The final dataset was collected on a Titan Krios (Thermo Fisher Scientific) operated at 300 kV and equipped with a K3 BioQuantum direct electron detector and energy filter (Gatan, Inc, AMETEK, Berwyn, PA, USA) using 20 eV slit. The data were acquired at 165 000 x nominal magnification with a calibrated pixel size of 0.504 Å. A total of 56 478 movies were collected in 33 frames with a total dose of 30 e^-^/Å2 (17 e^-^/pixel/s) over 0.45 s with a set defocus between -0.7 to -1.0 µm.

### Cryo-EM data processing

All processing was done on CryoSPARC v4.7.1-cuda12+250814 (29) following the workflow depicted in Supplementary Figure 1. Default parameters were used unless otherwise stated. The movies were motion-corrected using Patch Motion Correction, the contrast transfer functions (CTFs) were estimated using Patch CTF Estimation. Blob picker was used to produce an initial 3D reconstruction and produce templates for template picking. After template picking a particle stack was created by extracting a 800-pixel box around each particle binned to 600 pixels (0.672 Å/pix). 2D classification was used to identify obvious picking artifacts (*e.g*. carbon edges) and generate baits for sorting with heterogeneous refinement. Accepted and rejected particles from 2D classification were separately used to generate ab-initio models followed by heterogeneous refinement with all the particles. Classes corresponding to ribosomes were used in a homogeneous refinement performed simultaneously with higher-order CTF estimation (including spherical aberration, tetrafoil, anisotropic magnification) and per-particle defocus refinement, with Ewald’s sphere correction (negative sign) and estimation of per-particle scale factors. The resulting reconstruction was used for referenced-based motion correction and a new homogeneous refinement was performed with the same settings, resulting in a final consensus reconstruction with 1 823 753 particles. The 70S ribosome was subtracted using a mask that covered the ribosome and tRNA with a 20-pixel soft-edge (0.672 Å/pix). The resulting subtracted particles were then down-sampled to 128 pixels and a focused 3D classification was performed using a mask around the ribosomal A-site that covers the RRF•EF-G region with a 20-pixel soft-edge (0.672 Å/pix). The 3D classification was performed using the RRF•EF-G mask, keeping the input per-particle scales, filtering resolution of 6 Å, and RMS density convergence set to false. The number of 3D classes was iteratively increased to identify the optimum. For each job, the resulting classes were homogeneously refined using the original 600 pixel binned non-subtracted particles and the maps were examined to identify the job that produced classes with the best density for RRF and EF-G. The classification with 110 classes resulted in the highest-quality maps capturing the heterogeneity of the data. The particles containing RRF were additionally subjected to focused classification with a 40 Å radius spherical mask around the ribosomal A-site with 20 classes, keeping the input per-particle scales, filtering resolution of 6 Å, and RMS density convergence set to false. The particles in the classes containing RRF in the different conformations, EF-G, or RRF•EF-G were subjected to focused classification inside a SSU-mask (with 20-pixel soft edge) with 10 classes, while keeping the input per-particle scales. Homogeneous or non-uniform refinement using the original particles (600-pixel binned, 0.672 Å/pix) resulted in the final structures. Local-resolution estimation, Fourier shell correlation and angular distribution of the particles for the modeled structures are shown in Supplementary Figure 2. No 30S particles were identified during 2D or 3D classification.

All the final maps were post-processed using phenix.auto_sharpen (30) using the map and half-maps with local sharpening or low-pass filtered using RELION (31). The global resolution of maps was estimated within an auto-tightened mask created by CryoSPARC. For all maps used for model refinement, the resolution within a mask generated from the model was also calculated and used as the reported resolution, since the default auto-tightened masks from CryoSPARC did not cover the whole model in all the structures. These masks were created by calculating a synthetic map from the model using the molmap command in ChimeraX (32) at 20 Å, thresholded at a level where the map covers the whole model, and then adding a 20-pixel soft edge.

### Model building

The starting models used were 9GHG (33) (*S. aureus* 70S ribosome), 8P2G (22) (*S. aureus* EF-G) and an AlphaFold prediction of *S. aureus* RRF and *S. aureus* uL11 using the AlphaFold server (34). Each chain was first rigid-body fitted and refined using Coot (35) against the post-processed map, followed by global refinement using the sharpened maps using Servalcat (36). Afterwards, the structures were revised manually, and validation issues were corrected when possible. Validation was performed using Phenix (30). Refinement and validation information is found in Table 1.

**Table 1.** Cryo-EM refinement parameters and model validation.

|  | postTC•RRF•LSU<br>(EMDB-55945)<br>(PDB-9TI6) | postTC•RRF•LSU-2<br>(EMDB-55946)<br>(PDB-9TI7) | postTC•RRF•SSU<br>(EMDB-55944)<br>(PDB-9TI5) | 70S•RRF•EF-G•FA<br>(EMDB-55948)<br>(PDB-9TI9) | 50S•RRF•EF-G•FA<br>(EMDB-55947)<br>(PDB-9TI8) |
| --- | --- | --- | --- | --- | --- |
| **Data collection and processing** |  |  |  |  |  |
| Magnification | 165 000 | 165 000 | 165 000 | 165 000 | 165 000 |
| Voltage (kV) | 300 | 300 | 300 | 300 | 300 |
| Electron exposure (e<sup>-</sup>/Å<sup>2</sup>) | 30 | 30 | 30 | 30 | 30 |
| Defocus range (μm) | -0.7 to -1.0 | -0.7 to -1.0 | -0.7 to -1.0 | -0.7 to -1.0 | -0.7 to -1.0 |
| Pixel size (Å) | 0.504 | 0.504 | 0.504 | 0.504 | 0.504 |
| Symmetry imposed | C1 | C1 | C1 | C1 | C1 |
| Initial particle images (no.) | 2 556 350 | 2 556 350 | 2 556 350 | 2 556 350 | 2 556 350 |
| Final particle images (no.) | 25 462 | 24 569 | 123 631 | 9 322 | 77 333 |
| Map resolution (Å) | 2.62 | 2.76 | 2.20 | 3.06 | 2.18 |
| FSC threshold | 0.143 | 0.143 | 0.143 | 0.143 | 0.143 |
| **Refinement** |  |  |  |  |  |
| Initial model used (PDB code) | 9GHG, AlphaFold | 9GHG, AlphaFold | 9GHG, AlphaFold | 9GHG, AlphaFold, 8P2H | 9GHG, AlphaFold, 8P2H |
| Model resolution (Å) | 2.5 | 2.6 | 2.1 | 3.1 | 2.1 |
| FSC threshold | 0.5 | 0.5 | 0.5 | 0.5 | 0.5 |
| Post-processing | Phenix auto-sharpen | Phenix auto-sharpen | Phenix auto-sharpen | Phenix auto-sharpen | Phenix auto-sharpen |
| Map sharpening B factor. (Å<sup>2</sup>) | 15.87 | 14.73 | 6.65 | 48.03 | 15.02 |
| Model composition |  |  |  |  |  |
| Non-hydrogen atoms | 142 252 | 142 254 | 140 211 | 147 288 | 93 901 |
| Protein residues | 5 737 | 5 739 | 5 602 | 6 266 | 3 976 |
| RNA residues | 4 522 | 4 536 | 4 473 | 4 564 | 2 922 |
| Waters | 0 | 0 | 0 | 0 | 0 |
| Magnesium atoms | 122 | 113 | 127 | 90 | 115 |
| *B* factors (Å<sup>2</sup>) |  |  |  |  |  |
| Protein | 65 | 64 | 59 | 78 | 57 |
| RNA | 50 | 48 | 44 | 60 | 41 |
| Ligand | 26 | 23 | 26 | 35 | 38 |
| Water | - | - | - | - | - |
| R.m.s. deviations |  |  |  |  |  |
| Bond lengths (Å) | 0.008 | 0.008 | 0.008 | 0.008 | 0.008 |
| Bond angles (°) | 1.277 | 1.304 | 1.315 | 1.294 | 1.323 |
| Validation |  |  |  |  |  |
| MolProbity score | 1.00 | 1.13 | 1.03 | 1.14 | 0.96 |
| Clashscore | 1.41 | 2.15 | 1.80 | 2.68 | 1.93 |
| Poor rotamers (%) | 0.35 | 0.49 | 0.38 | 0.55 | 0.66 |
| Ramachandran plot |  |  |  |  |  |
| Favored (%) | 97.32 | 97.22 | 97.58 | 97.62 | 98.26 |
| Allowed (%) | 2.61 | 2.73 | 2.38 | 2.32 | 1.74 |
| Disallowed (%) | 0.07 | 0.05 | 0.04 | 0.06 | 0 |
| Rama-Z score | -1.41 | -1.26 | -1.03 | -1.16 | -0.68 |

SSU body rotation angles were calculated similarly as done by others (6). For SSU body rotation, the structures were superimposed by 23S rRNA and the angle between the axes formed by the 16S body residues (1-933 and 1409-1553) was calculated in ChimeraX. PDB entry 7NHM (37) was used as a classical-state reference. To assess the conformational change in RRF, the structures were aligned by domain I of RRF (1-29, 104-184) and the angle between the axes formed by domain II (30-103) was calculated in ChimeraX.

### Figures

Figures were rendered using UCSF ChimeraX, and assembled in Affinity Designer (Serif Europe Ltd, Nottinghamsire, United Kingdom).

## RESULTS

### Ribosome recycling captured by cryo-EM and fusidic acid inhibition

To study ribosome recycling, we assembled a postTC containing *S. aureus* 70S ribosomes, a short synthetic mRNA, deacylated *E. coli* tRNA^Ser^ and *S. aureus* RRF to which *S. aureus* EF-G, GTP and FA were added shortly before sample vitrification for cryo-EM analysis. Image processing identified 1 823 753 ribosomal particles, which were first subjected to 3D classification using a mask focused on the RRF and EF-G binding sites. This was followed by focused classification on the small subunit (SSU) to obtain homogenous structural states (Supplementary Figure 1). This resulted in several complexes containing RRF, EF-G or both solved at 2.2-3.1 Å global resolution (Figure 1, Supplementary Figure 2). The majority of ribosome particles corresponded to 70S postTC complexes carrying either EF-G locked by FA (33.8 %) or RRF alone (25.8 %). Notably, a small but well□resolved fraction of 70S particles contained both EF□G and RRF (1.1 %). Additional, less abundant complexes included 50S subunits bound to EF-G (2.7 %), RRF (3.6 %) or both factors (4.2 %). Because the 70S complexes with RRF displayed variability in SSU rotation, the highest-population states were modelled and analyzed in detail (Supplementary Figure 1). All complexes with EF-G showed density for FA.

**Figure 1:**
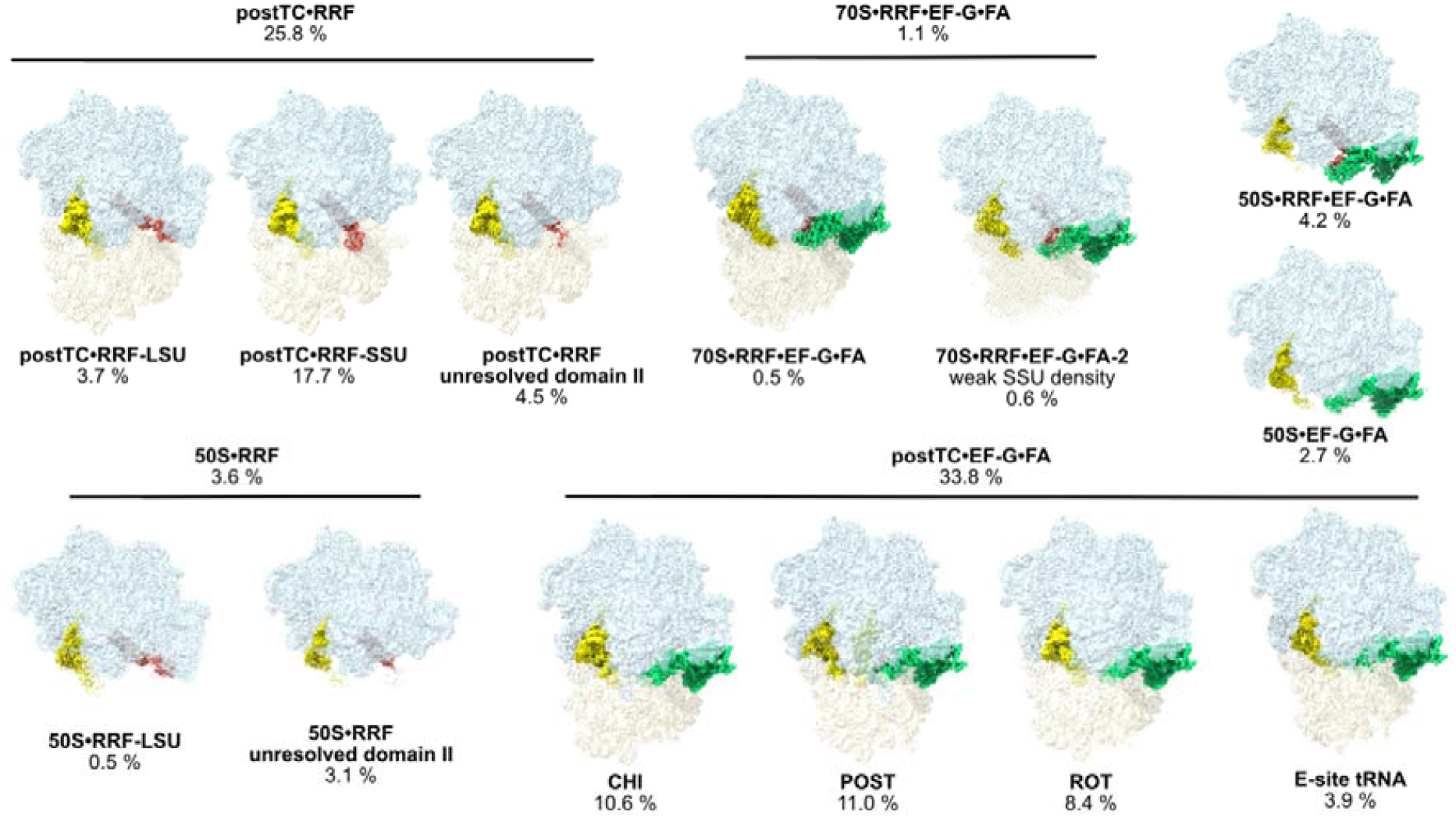
Cryo-EM analysis of FA inhibition of ribosome recycling. Cryo-EM reconstructions of ribosomal states containing RRF or EF-G identified in the dataset. Maps are segmented to show the LSU (light blue), SSU (light yellow), tRNA (yellow), EF-G (green), and RRF (salmon). The percentage of particles assigned to each state is indicated. States containing an empty A-site or A-site tRNA are included in Supplementary Figure 1.

These populations show that a considerably larger proportion of the 70S ribosomes are locked to only EF-G (33.8 %) than to both RRF and EF-G (1.1 %). Moreover, the majority of complexes harboring both factors are 50S subunits (4.2 %) rather than intact 70S ribosomes.

We observe EF-G trapped to the postTC in four main states with one or two tRNA molecules. The chimeric (CHI), rotated (ROT) and E-site states each have a single tRNA in either pe/E (CHI), p/E (ROT) or E-site conformation. The POST state contains two tRNAs in the P- and E-sites. The populations of the different states are comparable (Figure 1), with the state containing only E□site tRNA being the least abundant. We further found EF-G locked to 50S subunits (2.7 %), which likely reflects re□binding of EF□G to 50S rather than EF□G remaining locked after RRF departure.

### *S. aureus* RRF binds to the 70S ribosome in two main conformations

In agreement with previous observations (13), RRF binds to the postTC with domain I anchored to the large ribosomal subunit (LSU) and domain II in two distinct conformations. A main conformation shows domain II oriented towards the small ribosomal subunit (postTC•RRF-SSU, 17.7 %), while in a less abundant conformation faces the large ribosomal subunit (LSU) (postTC•RRF-LSU, 3.7 %) (Figure 1, 2 A-C, Supplementary Figure 1). In general, domain II of RRF appears more dynamic and reconstructs to lower resolution (Supplementary Figure 2) and in one set of particles domain II could not be clearly resolved (4.5 %, Figure 1).

**Figure 2:**
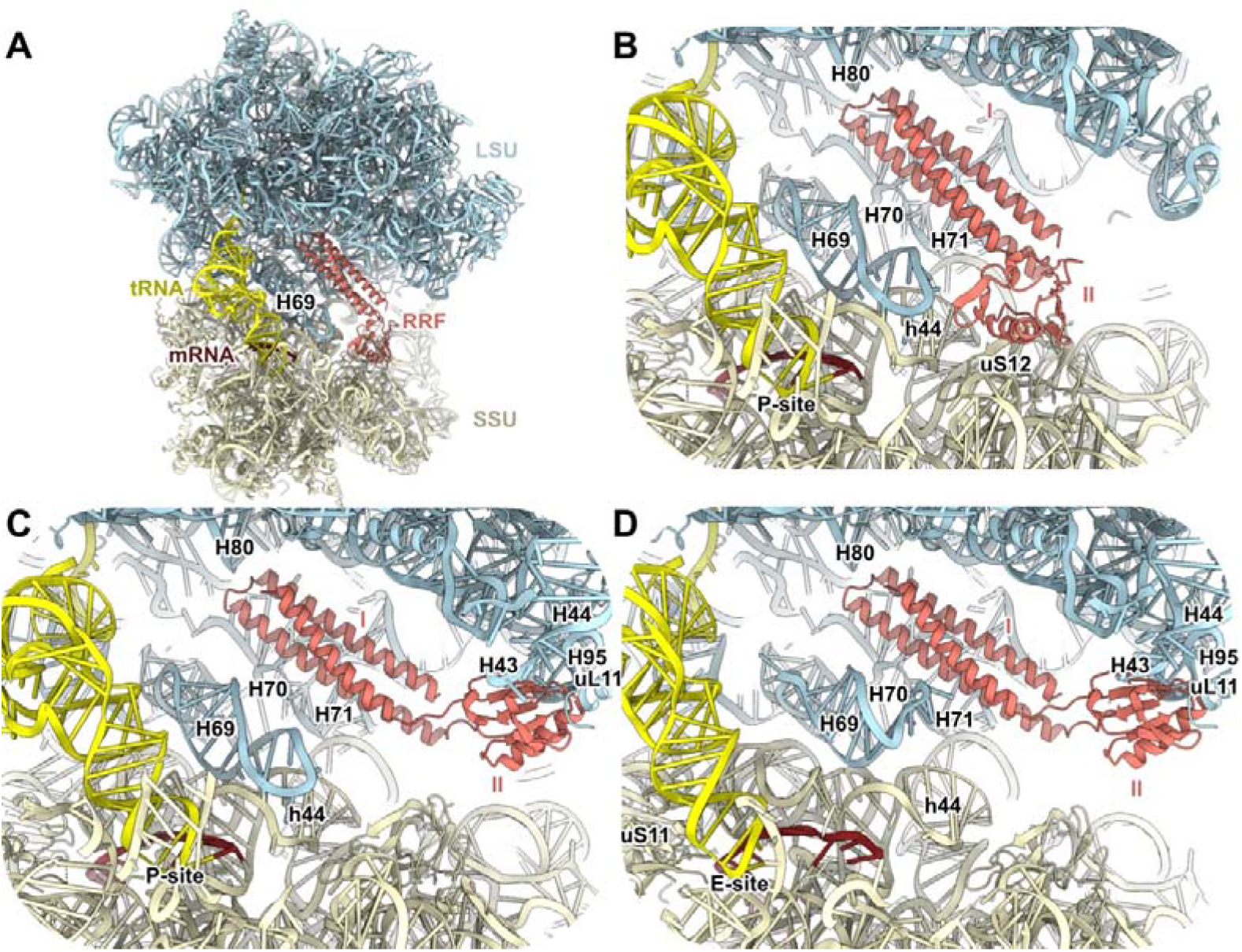
Structures of *S. aureus* RRF bound to the postTC. **(A)** Overview of the postTC•RRF-SSU structure showing the LSU (light blue), SSU (light yellow), tRNA (yellow), mRNA (brown) and RRF (salmon). **(B)** Enlarged view of (A) highlighting domain II of RRF contacting uS12. **(C)** Same view as in (B) for the postTC•RRF-LSU, in which domain II of RRF engages the LSU. **(D)** Same view as in (B) showing the postTC•RRF-LSU-2 state, with the tRNA positioned in the SSU E-site and H69 partially disengaged from h44.

In both conformations, domain I of RRF contacts only the LSU, interacting with the hairpin loop of H80 and extensively with H70 and H71 (Figure 2 B-C, Supplementary Figure 3 B-C). In the postTC•RRF-LSU conformation (Figure 2 C, Supplementary Figure 3 C), domain II extends towards the hairpin loops of H43, H44, and H95 (the sarcin-ricin loop, SRL) and uL11, without contacting the SSU. In contrast, in postTC•RRF-SSU, domain II of RRF contacts uS12 (Figure 2 A-B, Supplementary Figure 3 A-B). For both complexes, the most populated classes show the SSU in a rotated state and tRNA in a p/E hybrid state (Supplementary Figure 4). In the RRF-50S complexes, domain II is generally unresolved, but in a low-population class, domain II is observed interacting with the 50S subunit (Figure 1).

Interestingly, for postTC•RRF-LSU, one reconstruction is in close-to-classical state (1.8° body rotation and 1.4° swivel) and only carries an E-site tRNA (postTC•RRF-LSU-2, Figure 2 D, Supplementary Figures 3 D and 5 A). In this state, H69 is partially disengaged from the inter-subunit bridge B2a and instead interacts with domain I of RRF (Figure 2 D, Supplementary Figures 3 D and 5 B). This also occurs when EF-G is trapped on the ribosome with only an E-site tRNA (Supplementary Figure 5 C), suggesting that B2a is stabilized by P-site tRNA.

### Structures of RRF and EF-G bound to the 70S and 50S ribosome

We observe two well-resolved states of RRF and EF-G together on the ribosome: 50S•RRF•EF-G•FA and 70S•RRF•EF-G•FA. An additional, 70S•RRF•EF-G•FA-2 class was not interpreted due to low resolution and residual heterogeneity. Domain II of RRF is in all states oriented towards the SSU.

The 2.18 Å structure 50S•RRF•EF-G•FA allowed detailed interpretation of the interactions between RRF and EF-G on the ribosome (Figure 3, Supplementary Figure 6). As expected, FA is bound next to GDP and the SRL (Figure 3 B, Supplementary Figure 6 B). RRF mainly interacts with domains III-IV of EF-G through its domain II (Figure 3 C, Supplementary Figure 6 C), while domain I of RRF contacts domain III (Figure 3 D, Supplementary Figure 6 D) and the C-terminal helix of domain V of EF-G (Figure 3 E, Supplementary Figure 6 E). EF-G maintains its canonical contacts with the LSU at H43, H44 and H95.

**Figure 3:**
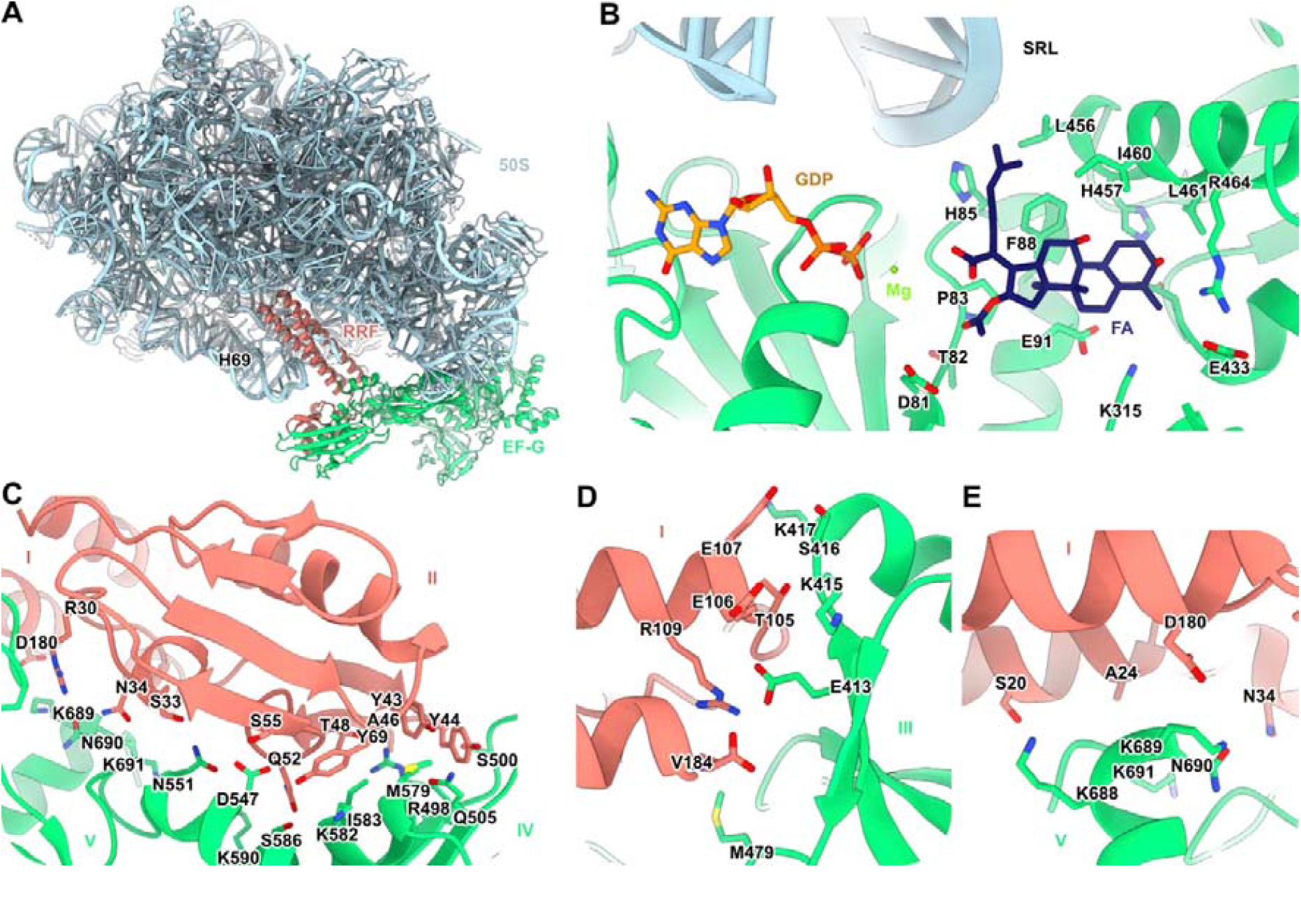
Structure of the 50S•RRF•EF-G•FA complex. **(A)** Structure overview, showing the LSU (light blue), RRF (salmon) and EF-G (green). **(B)** Close-up of the FA binding site showing EF-G residues within 4 Å of FA (dark blue), and GDP (orange) and magnesium ion (green). **(C)** Interaction between domain II of RRF and domain IV of EF-G. **(D)** Interaction between the C-terminus of RRF and domain II of EF-G. **(E)** Interaction between domain I of RRF and the C-terminus of EF-G.

The 70S•RRF•EF-G•FA state shows both RRF and EF-G bound to the 70S, indicating that this represents a ribosome recycling intermediate prior to subunit splitting (Figure 4 A). Although this reconstruction is of lower resolution (Supplementary Figure 2), the maps allowed confident modelling of each domain of RRF and EF-G (Supplementary Figures 7 A-B and 8), guided by the higher-resolution 50S•RRF•EF-G•FA structure. Notably, unlike during translocation, EF-G makes almost no contact with the SSU in this state (Figure 4 C).

**Figure 4:**
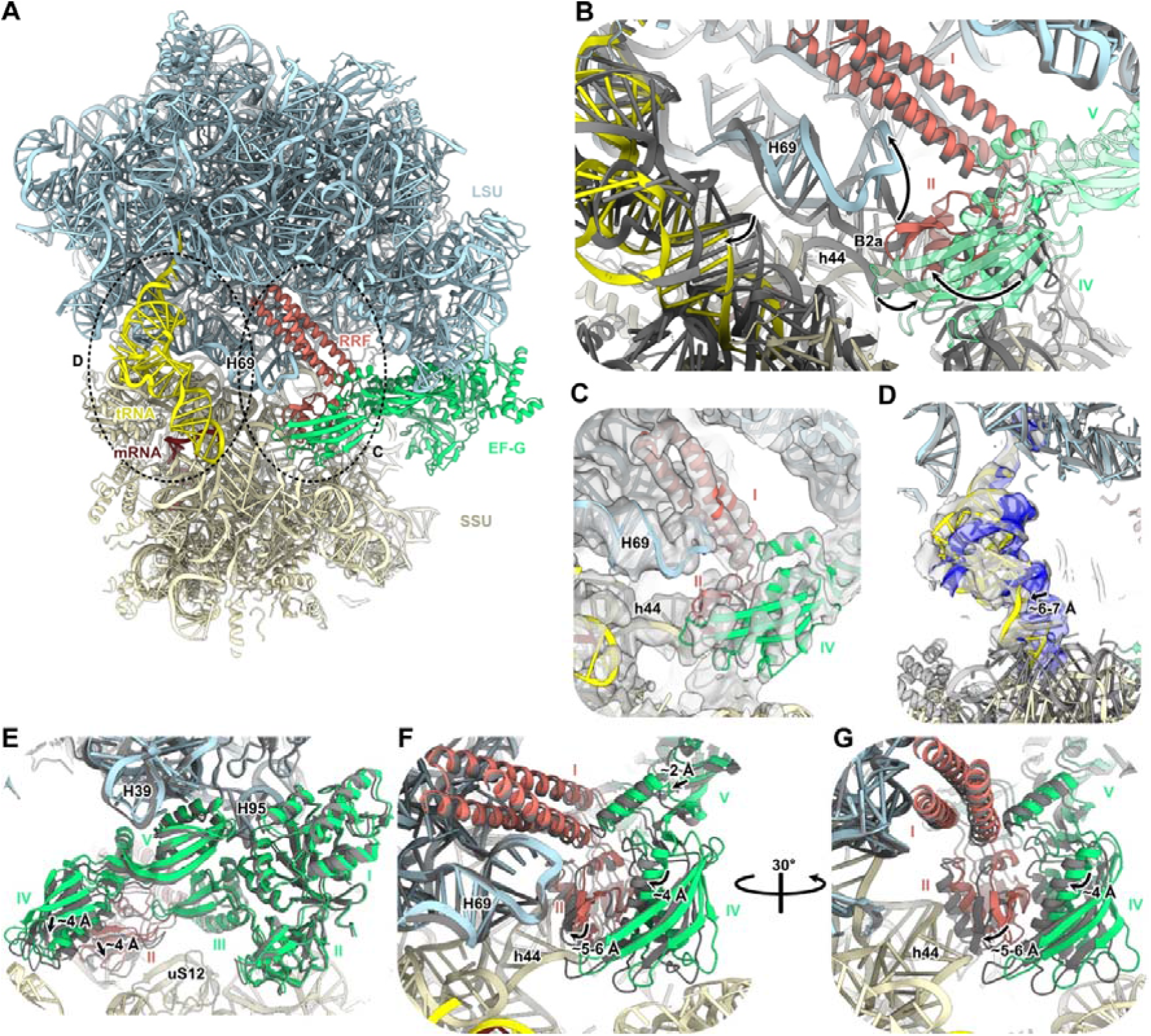
Structure of the 70S•RRF•EF-G•FA complex. **(A)** Structure overview, showing the LSU (light blue), SSU (light yellow), tRNA (yellow), mRNA (brown), RRF (salmon) and EF-G (green). **(B)** Conformational changes, indicated by arrows, between postTC•RRF-SSU (gray) and 70S•RRF•EF-G•FA (colored as in (A)) aligned by 23S rRNA. EF-G is shown semi-transparent. **(C)** H69 is disengaged from h44, breaking inter-subunit bridge B2a in 70S•RRF•EF-G•FA. The 6 Å low-pass filtered map is shown in gray. **(D)** The tRNA in 70S•RRF•EF-G•FA (gray map) is shifted from the P-site towards the SSU E-site relative to postTC•RRF-SSU (blue map). Maps are low-pass filtered to 6 Å. **(E-G)** Conformational changes, indicated by arrows, from 70S•RRF•EF-G•FA (colored as in (A)) to 50S•RRF•EF-G•FA (gray) aligned by 23S rRNA. **(E)** Conformational changes of EF-G. **(F-G)** Changes at the EF-G-RRF interface, showing how RRF domain II would clash with h44 of the 30S subunit.

### The combined action of RRF and EF-G destabilizes inter-subunit bridges and moves the tRNA from the P-site

The three RRF-bound structures along the recycling reaction pathway, postTC•RRF-SSU (EF-G substrate), 70S•RRF•EF-G•FA (recycling intermediate), and 50S•RRF•EF-G•FA (post-splitting product) allow direct comparison of the conformational changes during recycling. Binding of EF-G leads to several major structural changes (Figure 4 B, Supplementary Movie 1). In the 70S•RRF•EF-G•FA complex, domains III and IV of EF-G would sterically clash with domain II of RRF as positioned in postTC•RRF-SSU. Consequently, upon EF-G binding and GTP hydrolysis, domain II of RRF rotates by 32° relative to domain I, moving away from uS12 and towards inter-subunit bridge B2a, where it wedges between H69 and h44 (Figure 4 C), also causing rearrangement of h45. This action breaks bridge B2a, displacing H69 towards domain I of RRF and away from h44 of the SSU, which now interacts with domain II of RRF. These rearrangements collectively cause the SSU to back-rotate to a classical-like state (1.7 ° rotation, Supplementary Figure 7 C) and render the head dynamic (Supplementary Figure 2 E). Additionally, inter-subunit bridges B1b and B7b are broken, and bridges B6 and B8 are altered (Supplementary Figures 9-10). Simultaneously, the tRNA anticodon stem shifts approximately 6-7 Å from the SSU P-site towards the E-site (Figure 4 D). The density for the anticodon stem-loop is weaker than for the rest of the tRNA and surrounding rRNA (Supplementary Figure 7 D) and the mRNA density is only clearly identifiable for the Shine-Dalgarno (SD) sequence (Supplementary Figure 7 E). Thus, there is no clear support for preserved codon-anticodon interaction (Supplementary Figure 7 D-E). Although FA is present in both 70S•RRF•EF-G•FA states (Supplementary Figure 7 F-G), the low population of these complexes (1.1 %) compared to 50S•RRF•EF-G•FA (4.2 %) and postTC•EF-G•FA complexes (33.8 %) suggest that these are not the main FA-inhibited states. The observed loss of inter-subunit bridges and resulting SSU heterogeneity support the interpretation that this is a state on the path to subunit dissociation.

In the 50S•RRF•EF-G•FA complex, the interactions between RRF and EF-G are identical to in the 70S•RRF•EF-G•FA complex, and H69 remains in contact with RRF. However, both RRF and EF-G are shifted with respect to 23S rRNA (Figure 4 E-G). In particular, in 50S•RRF•EF-G•FA, EF-G domains IV-V and RRF domain II are positioned approximately 4-6 Å towards where the SSU would be (Figure 4 F-G), while domains I-III of EF-G and domain I of RRF are situated similarly in both structures (Figure 4 E). This shows that RRF and EF-G in presence of fusidic acid adopt a structure on the 50S subunit that would be incompatible with a bound 30S subunit.

There is weak density for the E-site tRNA (Figure 1), possibly due to rebinding. The high similarity, including contacts with H69, suggests that this complex is the product of ribosome splitting in presence of FA. However, we cannot exclude that this complex is a product of binding and FA-locking of free EF-G to the 50S•RRF complex.

Interestingly, some of the inter-subunit bridges are to a smaller extent perturbed in absence of EF-G and P-site tRNA in postTC•RRF-LSU-2 (Supplementary Figures 9-10), where bridge B2a is partially broken and B7b is absent. This showcases the relevance of destabilizing the tRNA-SSU interaction during recycling (Figure 4 D, Supplementary Figure 7 D), as the P-site tRNA connects the head and body of the SSU with the LSU.

## DISCUSSION

In this study, we have used cryo-EM to visualize the inhibition of ribosome recycling by FA in the clinically relevant bacterium *S. aureus*. Previous cryo-EM structures of RRF and EF-G bound together to postTCs or 50S subunits (13, 38, 39) were limited by resolution and 3D classification methodology. Here, we have captured a previously unobserved late intermediate state of ribosome recycling prior to subunit splitting (70S•RRF•EF-G•FA) and obtained high-resolution maps of an RRF-EF-G-50S complex. These structures provide mechanistic detail of how the sequential binding of RRF and EF-G leads to ribosome splitting, and how FA inhibits this process.

Several studies have demonstrated the co-evolution of EF-G and RRF to ensure functional interactions for recycling (40–42). Comparison of our 70S•RRF•EF-G•FA complex with a heterologous postTC with *T. thermophilus* RRF, *E. coli* EF-G and FA (39) explains why TtRRF and *Ec*EF-G are incompatible and not splitting competent (40). In the structure of the heterologous complex, domain II of *Tt*RRF is positioned between domains IV and V, on the opposite side of domain IV of *Ec*EF-G, instead of approaching bridge B2a. The interactions observed in our structure also do not agree with the crystal structure of the 70S ribosome with RRF and an EF-G-L9 fusion, where EF-G presents a compact conformation with domain IV oriented away from RRF (43).

Based on time-resolved cryo-EM structures, it was hypothesized that EF-G would help domain II of RRF to attack inter-subunit bridge B2a (13, 39). This was supported by experiments showing that ribosomes lacking H69 can be recycled in the absence of RRF (44). However, there was no direct observation of the process of subunit separation, and the density for domain II of RRF appeared fused to that of EF-G. Thus, our 70S•RRF•EF-G•FA structure, presenting clear density for how RRF domain II wedges apart bridge B2a and breaks inter-subunit contacts (Figure 4, Supplementary Figures 9-10) confirms the initial hypothesis. Furthermore, it had been unclear whether ribosome recycling involved back-rotation of the SSU. Our structures show that the rearrangement in h44 is associated with back-rotation of the SSU, disruption of bridges B1b and B7b, changes of B6 and B8, and destabilization of the tRNA. Ribosome splitting by EF-G and RRF is faster in presence of deacylated P-site tRNA than for vacant ribosomes. This is consistent with previous findings that deacylated tRNA favors the rotated ribosome conformation (11), required for binding of RRF and EF-G (45).

A long-standing question is how tRNA is released during recycling (20, 45–47). Kinetic experiments have demonstrated a correlation between the SD-P-site codon distance and the binding stability of deacylated P-site tRNA (47). In absence of an upstream SD, it was shown that mRNA dissociates first, then tRNA leaves and finally splitting occurs, while for SD-containing mRNAs, the three events appear to coincide (20). In our sample, containing a strong SD sequence, the 70S•RRF•EF-G•FA structure shows that the tRNA anticodon has lost contact with the P-site in the SSU (Figure 4 D, Supplementary Figure 7 D), consistent with tRNA dissociation upon splitting, while the mRNA remains bound to the SSU via SD interactions (Supplementary Figure 7 E). In the absence of an SD sequence, it is possible that the mRNA would dissociate from the 30S. Ribosome splitting is 3-fold slower with tRNA^fMet^ than with tRNA^Phe^ (48). This can be explained by the interactions between the conserved G-C base pairs of initiator tRNAs and the SSU P site (49), which according to our structure, would break during recycling. In support of the requirement of displacement of the P-site tRNA to break B2a, previous 70S•RRF structures that observed changes in H69 (15, 17) all lacked P-site tRNA, as in postTC•RRF-LSU-2. However, there is no translocation of mRNA during recycling (47, 50), showing that the disordered anticodon does not pull along the mRNA.

With these new structural insights, we can propose a model for the mechanism of ribosome recycling and FA inhibition (Figure 5). Following termination, a deacylated tRNA remains paired to the P-site mRNA codon on the 70S ribosome (postTC). This state allows SSU rotation, bringing the tRNA into the p/E site. RRF binds and stabilizes the rotated ribosome (45, 48), with domain I fixed by LSU interactions and domain II adopting multiple orientations (Figure 2). In the most populated state, domain II is positioned towards the SSU (postTC•RRF-SSU). Only this state is sterically compatible with initial binding of EF-G•GTP, quickly triggering GTP hydrolysis (18). In the resulting complex, domains III-IV of EF-G push RRF towards inter-subunit bridge B2a, where RRF interactions cause rearrangement of H69 and h44, breaking bridge B2a (Figure 4 B-C). In addition, conformational changes of h44 and h45 cause the SSU to back-rotate (Supplementary Figure 7 C), shifting the tRNA towards the 30S E-site (Figure 4 D) and de-stabilization of the anticodon stem-loop (Supplementary Figure 7 D).

**Figure 5:**
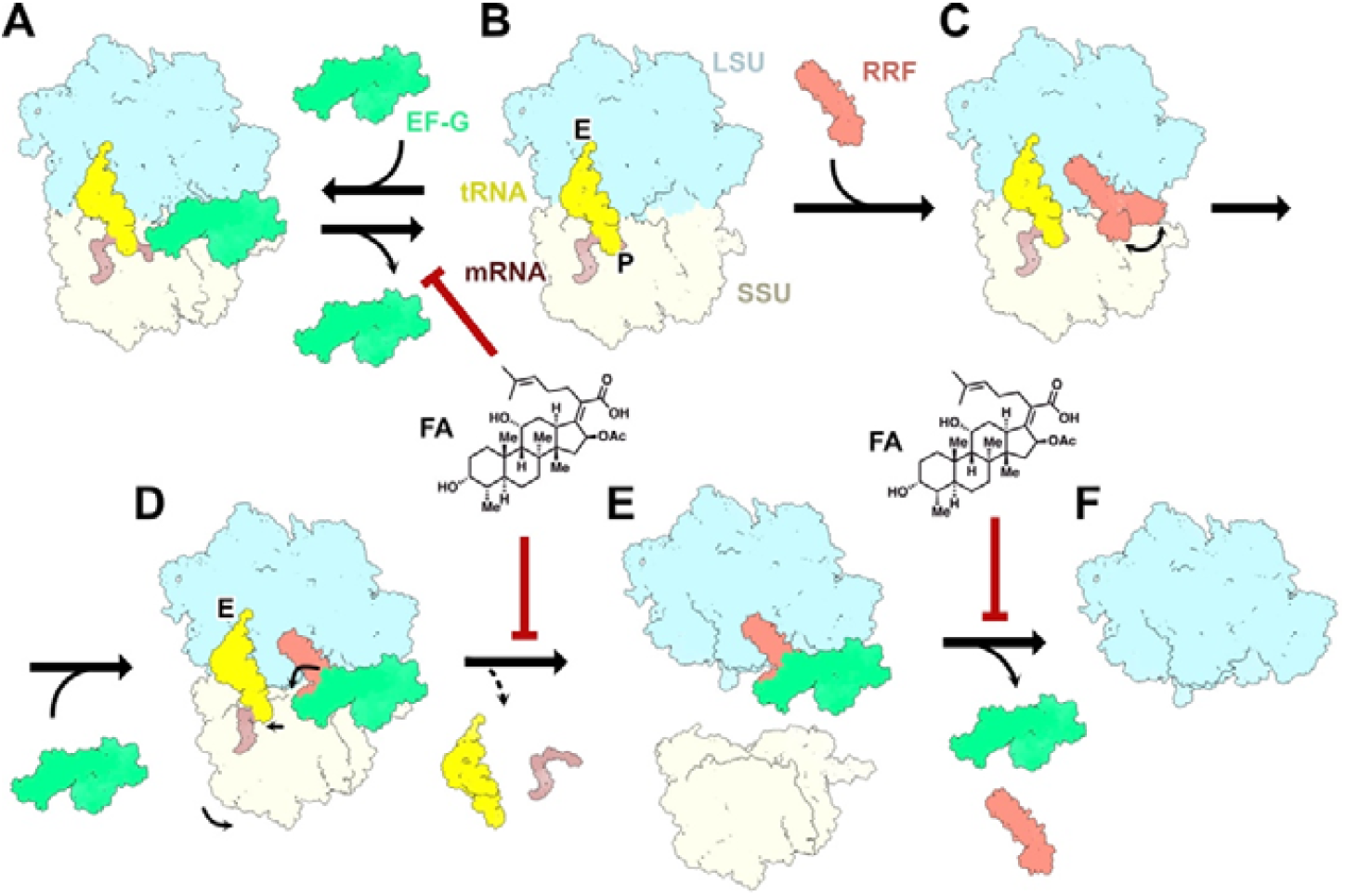
Mechanism of ribosome recycling and inhibition by FA. The postTC comprises the ribosome (LSU in light blue and SSU in light yellow), deacylated tRNA (yellow) and mRNA (brown). **(A)** EF-G (green) binds to the postTC, hydrolyzes GTP and dissociates. The dissociation is inhibited by FA. **(B)** postTC. (C) Upon binding to postTC, RRF (salmon) adopts multiple orientations of domain II. **(D-E)** EF-G binds to the RRF-bound postTC. Their combined action induces SSU back-rotation, disrupts inter-subunit bridges and destabilizes tRNA interactions with the SSU, leading to splitting of the 70S ribosome. In absence of a nearby SD-sequence, tRNA and mRNA are released. FA inhibits splitting, but not tRNA and mRNA release. **(E-F)** Release of RRF and EF-G from the 50S subunit. The dissociation is inhibited by FA.

The movement propagates to bridges B1a, B1b, B7b, B6, and B8, which are broken or altered (Supplementary Figures 9-10), consistent with subunit splitting and tRNA release. This would be followed by departure of EF-G•GDP from the 50S, followed by dissociation of RRF.

The 70S•RRF•EF-G•FA complex is considerably less abundant than other EF-G•FA complexes, and shows multiple disrupted inter-subunit bridges (Supplementary Figures 9-10), as well as limited contacts between EF-G and the SSU. According to biochemical studies with *E. coli* components, FA inhibits subunit splitting (20, 26), but not release of tRNA and mRNA (20). Thus, the 70S•RRF•EF-G•FA complex is expected to remain stable while FA is bound; on the order of ∼5 seconds, but is “primed” for ribosome splitting upon FA dissociation (26). The observed wedging of RRF domain II into bridge B2a and the movement of the P-site anticodon stem out of the SSU P-site supports that 70S•RRF•EF-G•FA represents an on-path intermediate of the recycling reaction, and is consistent with splitting upon FA dissociation. The 50S•RRF•EF-G•FA structure shows an additional shift of RRF domain II that would severely clash with the SSU (Figure 4 F-G), suggesting that a similar movement in absence of FA would complete subunit splitting. In this model, the abundant 50S•RRF•EF-G•FA complex could thus be a result of FA rebinding after subunit splitting but before EF-G dissociation from 50S•RRF.

Our structural data would also be compatible with a model where the *S. aureus* 70S•RRF•EF-G•FA complex is splitting-competent, and that RRF and EF-G would remain trapped on the resulting 50S product, but this would be mechanistically different from the *E. coli* system.

However, our structures and population distribution support a model in which FA does not primarily inhibit ribosome recycling by preventing splitting of postTCs bound to RRF and EF-G. Instead, both models support that FA primarily inhibits recycling by trapping EF-G to the postTC in absence of RRF, thereby preventing RRF binding and blocking the splitting reaction (26). Measurements of GTP hydrolysis by EF-G per postTC splitting event at varying RRF concentrations showed that, under saturating concentration of RRF, the fraction of non-productive GTP hydrolysis cycles not leading to subunit splitting is negligible (18). Further, the GTP consumption per ribosome splitting event is not affected by FA (26). Based on concentrations of RRF and EF-G in *E. coli*, it was estimated that five GTP molecules would be consumed per ribosome-splitting event *in vivo* (18). This suggests that the majority of postTC-EF-G complexes available for FA binding lack RRF and that for recycling, this mode of inhibition may also be predominant *in vivo*.

During translocation, GTP hydrolysis and Pi release from EF-G unlocks the decoding center and brings the ribosome into a chimeric state of the SSU, through back-rotation and head swivel (6, 7). These changes involve direct interactions between EF-G and the SSU. During recycling, on the other hand, EF-G also promotes SSU back-rotation, but mediated by RRF, which in addition leads to disruption of inter-subunit bridges. This results in a state with very limited contacts between EF-G and the SSU, but extensive EF-G-RRF contacts. The requirement of GTP hydrolysis for ribosome splitting was biochemically demonstrated (1, 50) and later confirmed by single-molecule experiments with the non-hydrolysable GTP analogue GDPNP (45). This indicates that EF-G-GDPNP is incapable of changing the conformation of RRF (47), inhibiting the reaction at an earlier state than observed with FA, which binds after Pi release. Interestingly, the earliest-state structures of EF-G during translocation, with tRNA in A/P state, prior to Pi release (6), would severely clash with postTC•RRF-SSU (Supplementary Figure 11 A), and the GDP state of EF-G would clash with RRF in 70S•RRF•EF-G•FA (Supplementary Figure 11 B). This indicates that at initial binding, prior to Pi release, as well as at the end of the reaction, the conformation of EF-G must be different during recycling compared to during translocation. However, time-resolved cryo-EM suggests that EF-G domains I-II dissociate first from the ribosome after translocation, leading to small conformational changes in EF-G domains III-V (6). A similar sequence of events might occur during recycling, and the conformational state of domains IV-V observed in the 50S•RRF•EF-G•FA structure may correspond to this later stage.

In summary, this study captured FA inhibition of ribosome recycling in *S. aureus*, visualizing intermediates in which inter-subunit bridges are disrupted by RRF•EF-G, leading to SSU back-rotation, tRNA movement and, eventually, subunit splitting. The data supports a model in which FA primarily inhibits recycling by trapping EF-G to the postTC in absence of RRF rather than preventing EF-G from completing RRF-mediated ribosome splitting. Together, these findings provide structural detail for RRF•EF-G-mediated recycling and its inhibition by FA.

## ACKNOWLEDGEMENTS

We thank Måns Ehrenberg and Anneli Borg for critical discussion of the results. We thank Daniel Larsson for cryo-EM data collection, discussions and comments on the manuscript and Shirin Akbar for comments on the manuscript. We acknowledge the use of the Cryo-EM Uppsala facility for grid preparation and screening, funded by the Department of Cell and Molecular Biology, the Disciplinary Domains of Science and Technology and of Medicine and Pharmacy at Uppsala University. We acknowledge access and support of the cryo-EM facilities at the UK national electron Bio-Imaging Centre, proposal BI42543.

## AUTHOR CONTRIBUTIONS

Adrián González-López: conceived the project, designed experiments, purified the ribosomes and proteins, prepared cryo-EM samples, collected and processed cryo-EM data, modeled structures, analyzed structures and wrote the manuscript. Maria Selmer: conceived the project, designed experiments, analyzed structures, wrote the manuscript and secured funding.

## DATA AVAILABILITY

The cryo-EM maps and models in this study have been deposited in the Electron Microscopy Data Bank under the accession codes 9TI5 (postTC•RRF-SSU), 9TI6 (postTC•RRF-LSU), 9TI7 (postTC•RRF-LSU-2), 9TI8 (50S•RRF•EF-G•FA), and 9TI9 (70S•RRF•EF-G•FA), and Protein Data Bank under the accession codes EMD-55944 (postTC•RRF-SSU), EMD-55945 (postTC•RRF-LSU), EMD-55946 (postTC•RRF-LSU-2), EMD-55947 (50S•RRF•EF-G•FA), and EMD-55948 (70S•RRF•EF-G•FA). Raw cryo-EM movies, particle stacks and extra maps have been deposited in the Electron Microscopy Image Archive under the accession code EMPIAR-13144. Custom python scripts used during modeling and analysis are available on Git Hub (https://github.com/adriangl97/pdb_python_tools).

