## Supplementary material for "Structural characterization of ribosome recycling and fusidic acid inhibition in *Staphylococcus aureus*"

**Content:**

**Supplementary Figures 1-11**

**Supplementary references**

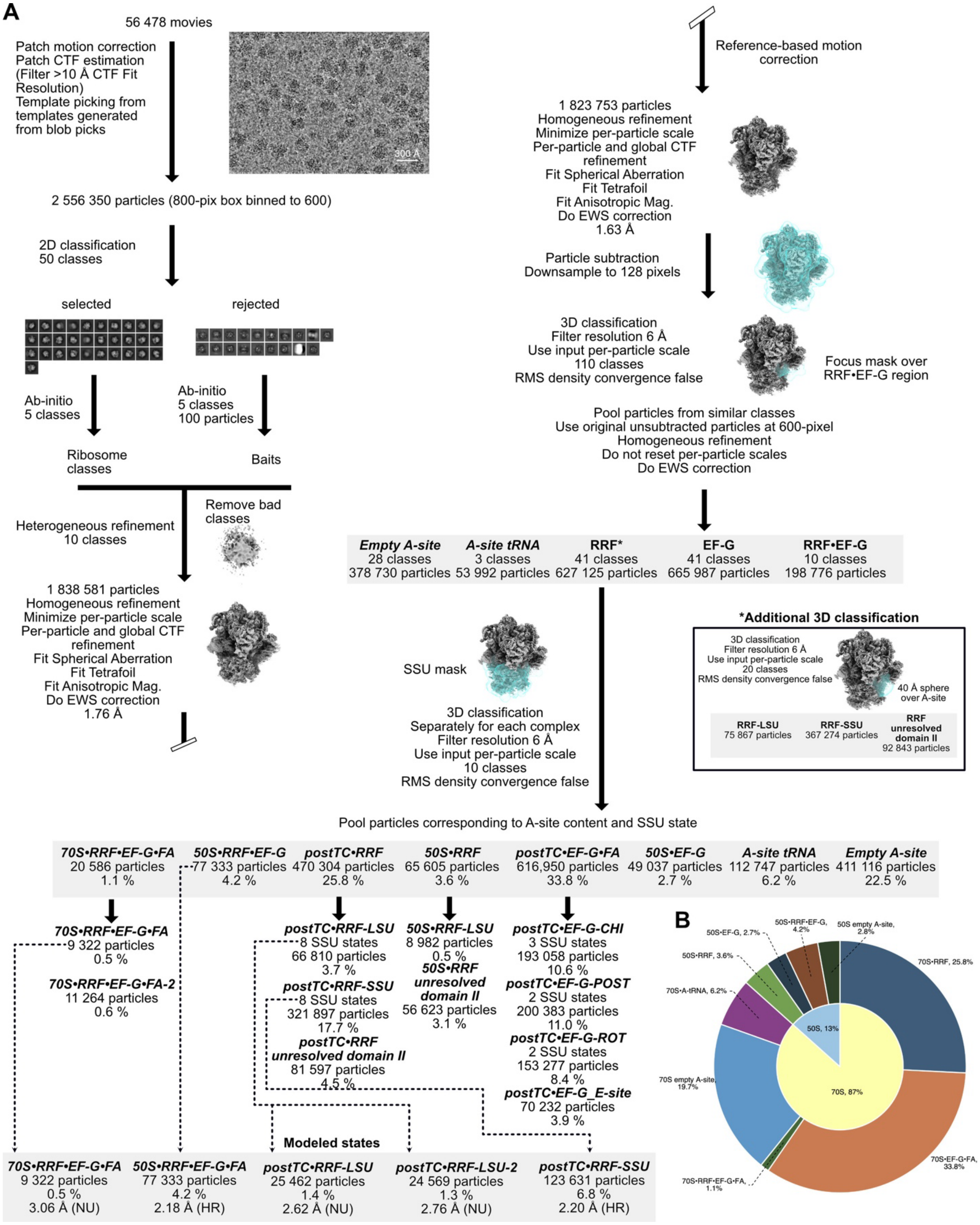


**Supplementary Figure 1: Processing workflow using CryoSPARC (1).** SSU states, particle numbers and their percentage of total ribosomes are indicated in text **(A)** and a pie chart **(B)**.

For states subjected to modelling, the final resolution and refinement algorithm (NU, non-uniform refinement or HR, homogenous refinement) are reported.

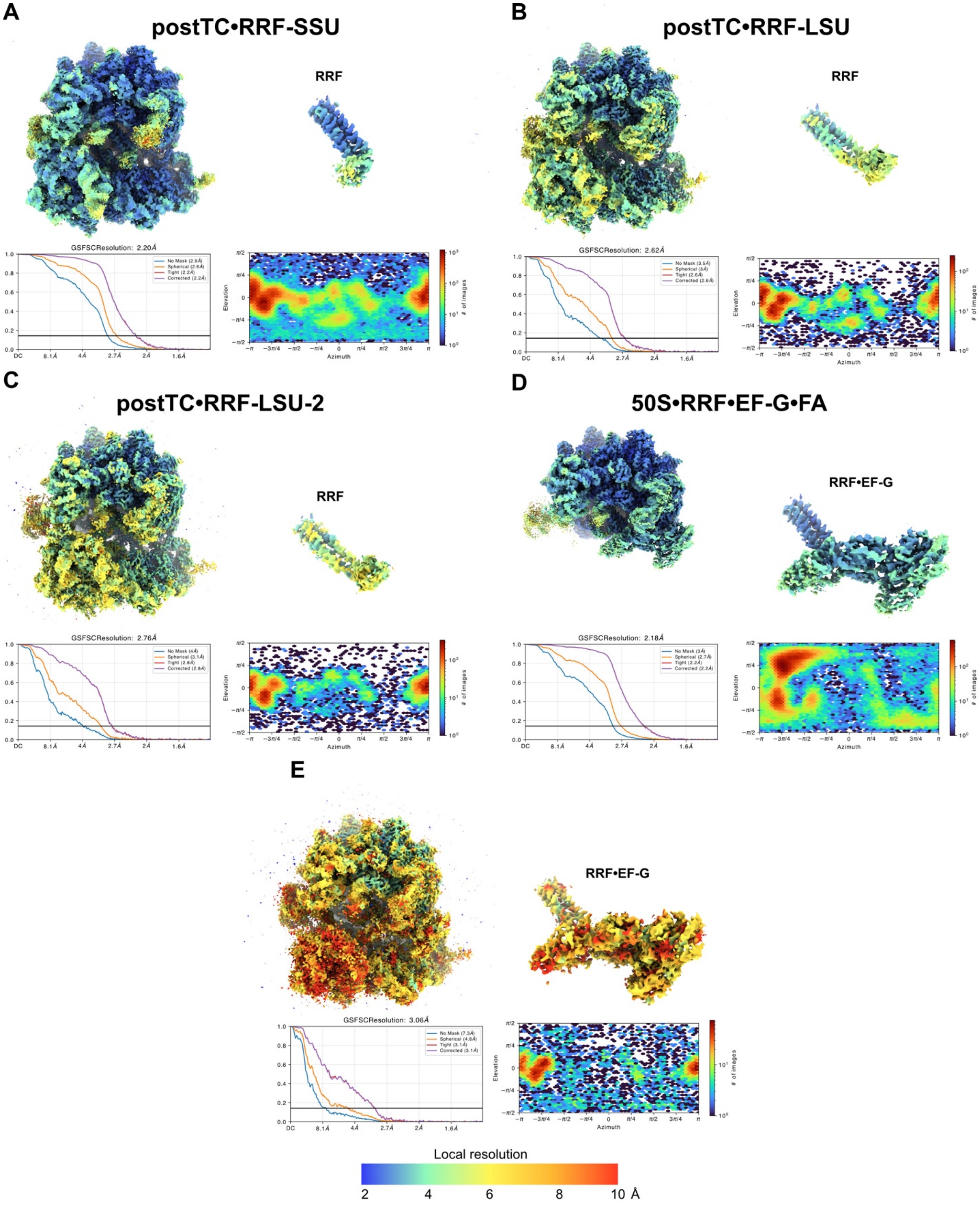


**Supplementary Figure 2: Resolution assessment of modeled cryo-EM reconstructions.** For each reconstruction, un-sharpened maps and segmented densities within 3 Å of the models highlight RRF or RRF•EF-G are colored by local resolution estimates from CryoSPARC (top). Global Fourier shell correlation curves and particle angular distributions (bottom). **(A)** postTC•RRF-SSU. **(B)** postTC•RRF-LSU. **(C)** postTC•RRF-LSU-2. **(D)** 50S•RRF•EF-G•FA. **(E)** 70S•RRF•EF-G•FA.

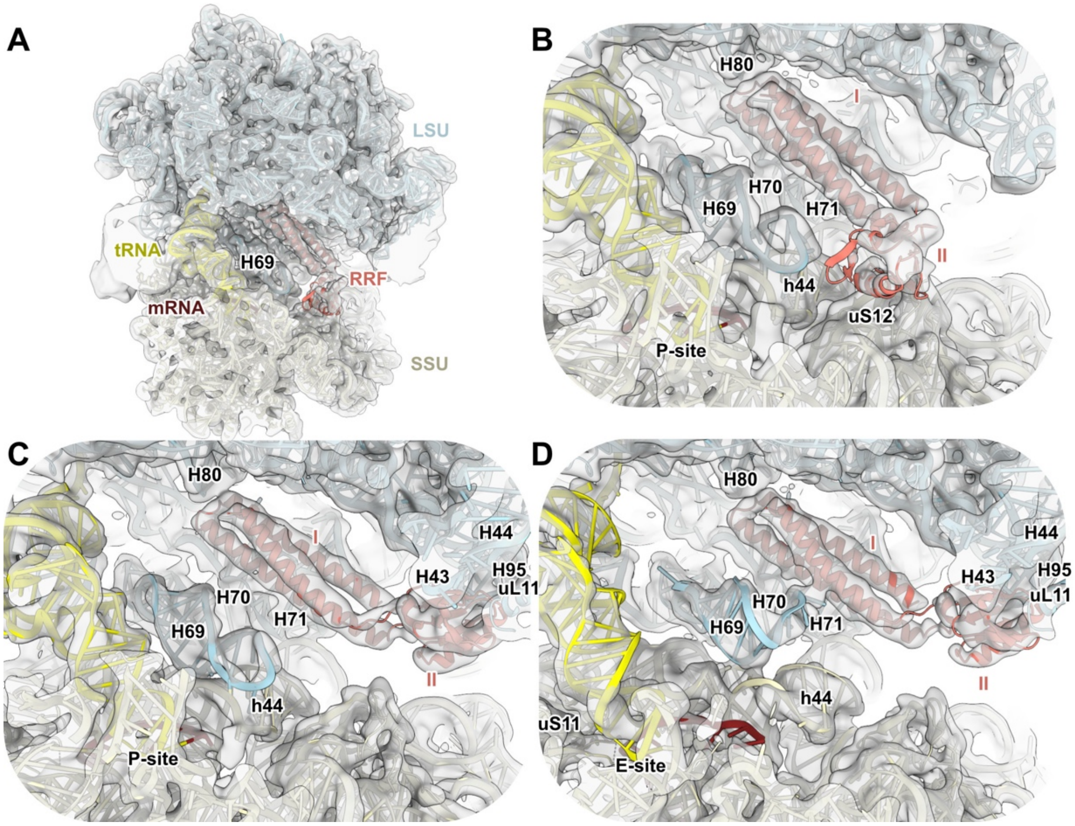


**Supplementary Figure 3: Locally sharpened maps around regions shown in Figure 2.** As Figure 2 but showing 6 Å low-pass filtered maps (gray) for all states.

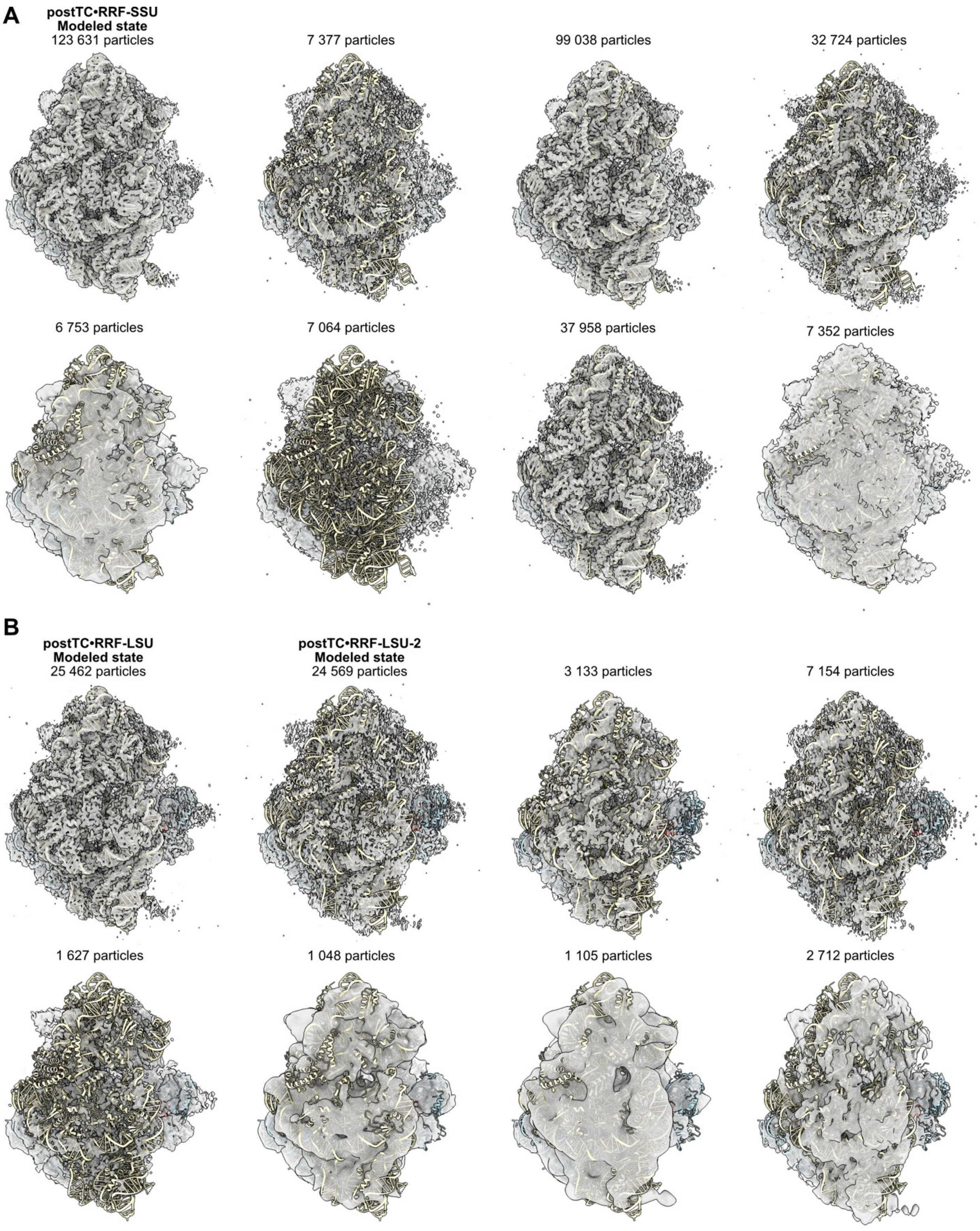


**Supplementary Figure 4: SSU states of 70S particles containing RRF.** Structures are shown from the SSU solvent side with LSU in light blue, SSU in light yellow and maps are displayed in gray. Particle numbers and whether each state was modeled are indicated. **(A)** postTC•RRF-SSU. **(B)** postTC•RRF-LSU.

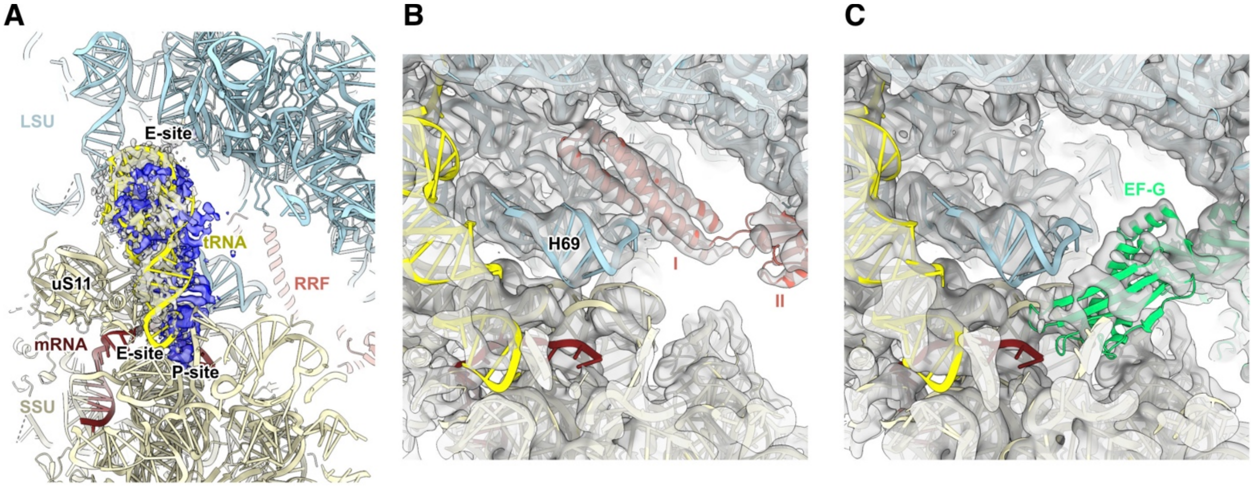


**Supplementary Figure 5: The postTC•RRF-LSU-2 structure and effects of P-site tRNA absence. (A)** Position of the tRNA (yellow) in postTC•RRF-LSU-2. LSU (light blue), SSU (light yellow), mRNA (brown) and RRF (salmon) are shown. Locally sharpened maps from postTC•RRF-LSU-2 (gray) and postTC•RRF-LSU (blue) are segmented within 3 Å of the tRNA in their respective models. **(B)** 6 Å low-pass filtered map of postTC•RRF-LSU-2 around H69. **(C)** 6 Å low-pass filtered map of postTC•EF-G with an E-site tRNA. The models shown are postTC•RRF-LSU-2 without RRF, and EF-G (green) from PDB 8P2G (2), rigid-body fitted into the map for illustration purpose**.**

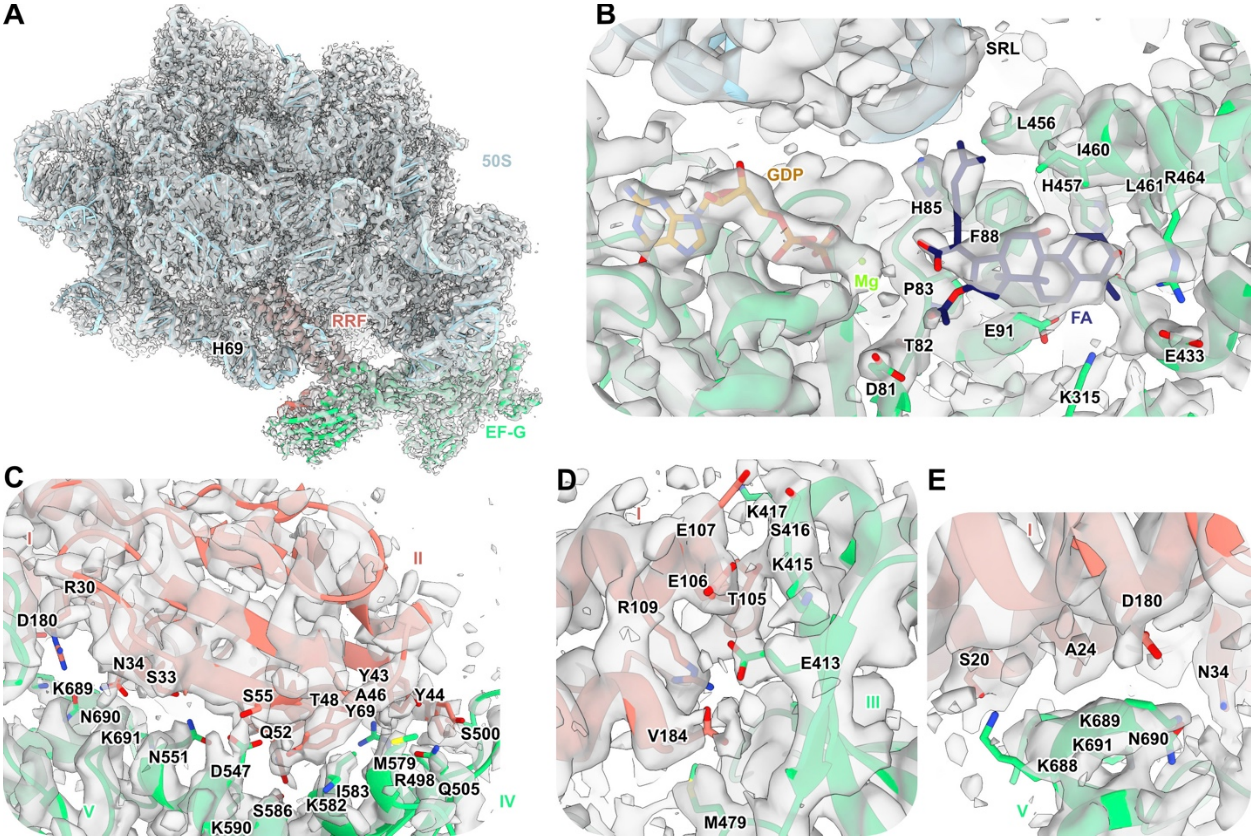


**Supplementary Figure 6: Locally sharpened maps around regions shown in Figure 3.** As Figure 3 but showing locally sharpened maps (gray). In (A), the map is segmented within 3 Å of the model.

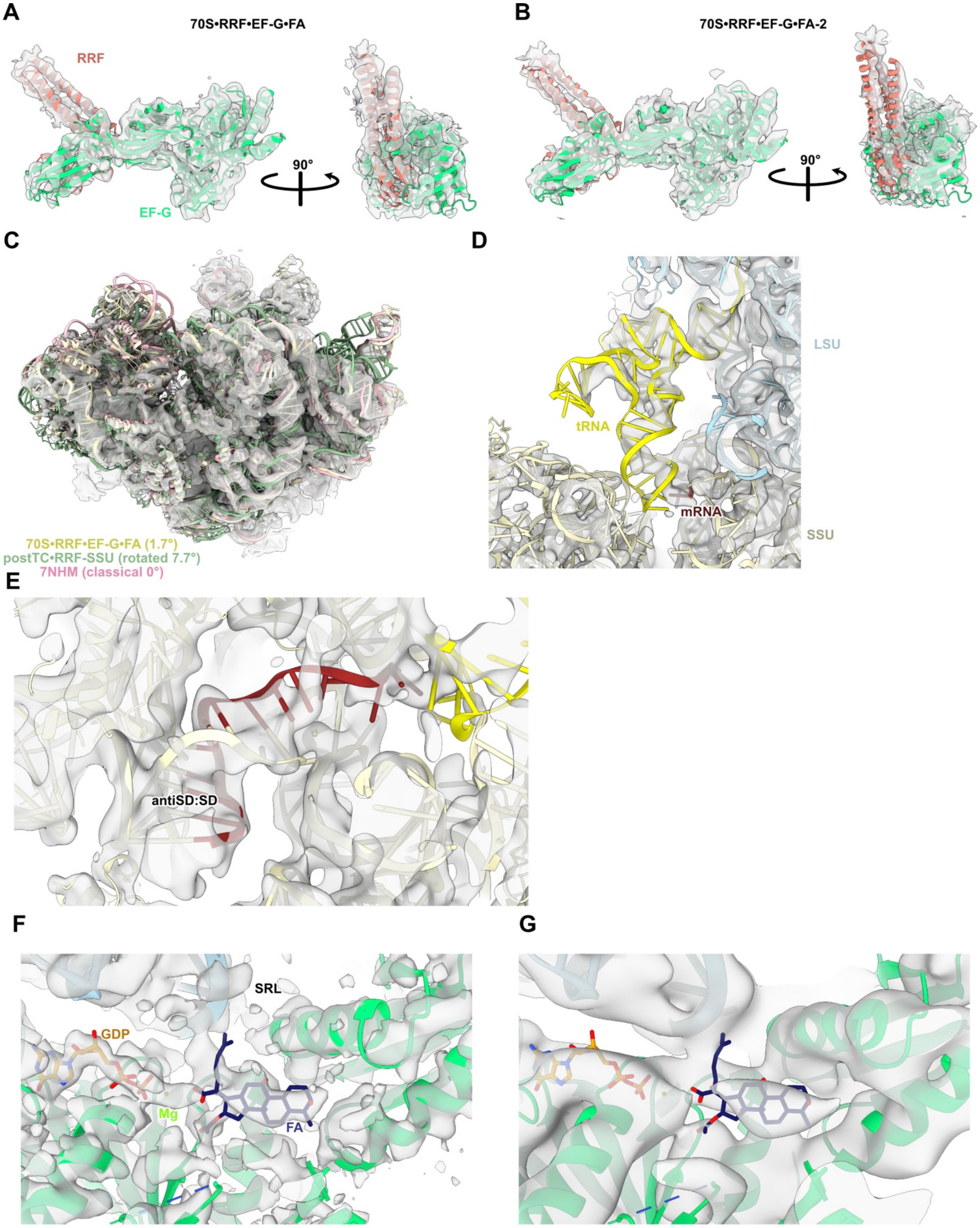


**Supplementary Figure 7: Cryo-EM maps of 70S•RRF•EF-G•FA. (A)** Segmented density within 3Å of RRF•EF-G (salmon and green) from the 6Å low-pass filtered 70S•RRF•EF-G•FA map (gray).

**(B)** Same as (A) for the 70S•RRF•EF-G•FA-2 map. **(C)** SSU solvent-side of 70S•RRF•EF-G•FA (SSU, light yellow; LSU, light blue; EF-G, green) with 6 Å low-pass filtered map (gray). Structures of postTC•RRF-SSU (light green, rotated state) and *S. aureus* 70S in classical state (pink, PDB 7NHM(3)) are superimposed based on 23S rRNA. **(D)** 6 Å low-pass filtered map of 70S•RRF•EF-G•FA (gray) around the tRNA (yellow). LSU (light blue), SSU (light yellow), mRNA (brown) are shown. **(E)** 6 Å low-pass filtered of 70S•RRF•EF-G•FA (gray) around the SD:antiSD helix. Colors as in (c). **(F)** Local sharpened map of 70S•RRF•EF-G•FA (gray) at the FA binding site showing FA (dark blue), GDP (orange) and a magnesium ion (green). **(G)** Same region as in (E) but for the 6 Å low-pass filtered map of 70S•RRF•EF-G•FA-2.

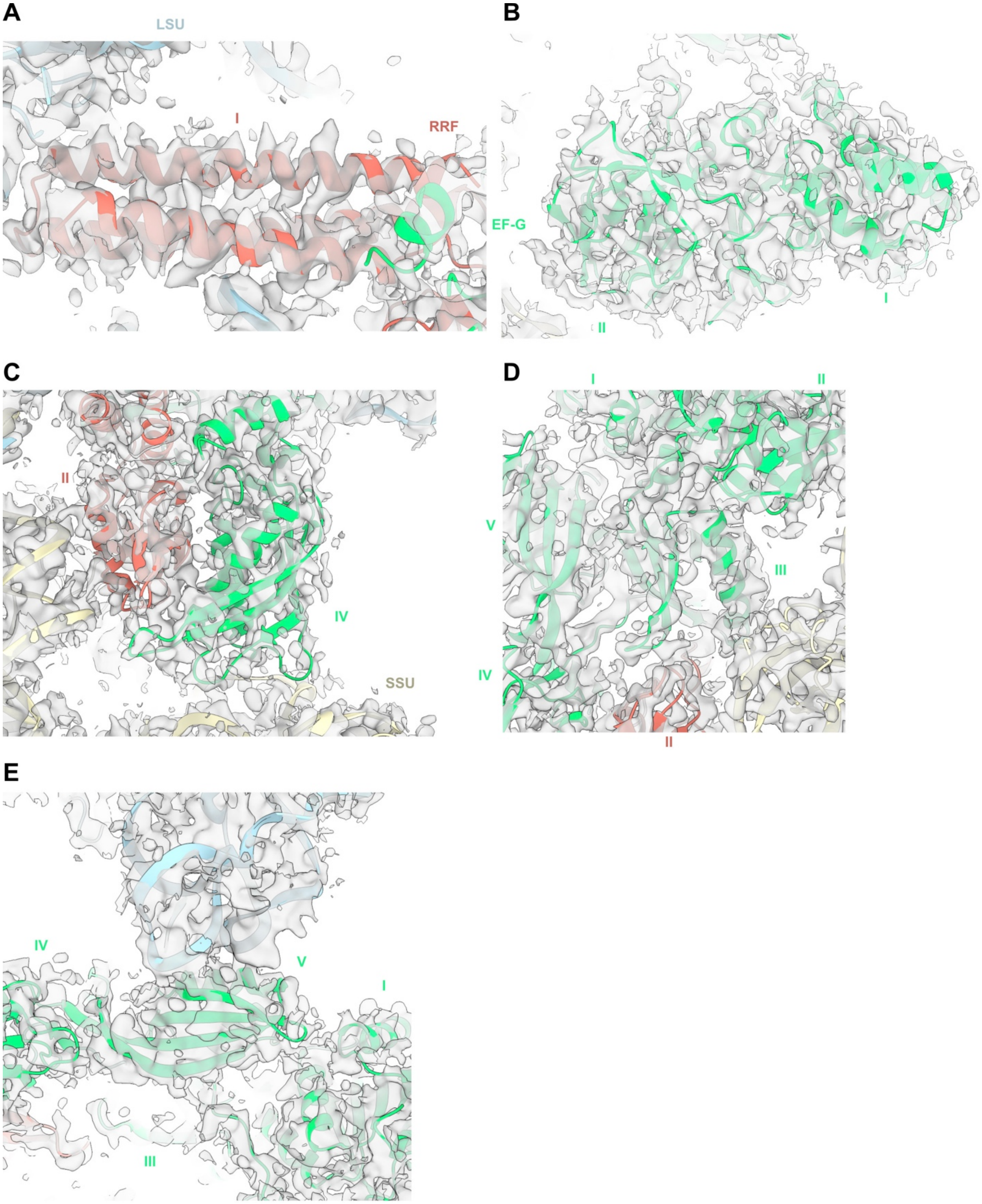


**Supplementary Figure 8: Locally sharpened map of 70S•RRF•EF-G•FA.** Locally sharpened map (gray) segmented 3 Å around the 70S•RRF•EF-G•FA model. LSU (light blue), SSU (light yellow), RRF (salmon) and EF-G (green) are displayed. **(A)** RRF domain I. **(B)** EF-G domains I-II.

**(C)** RRF domain II with EF-G domain IV. **(D)** RRF domain II with EF-G domains I-V. **(E)** EF-G domains I and III-V.

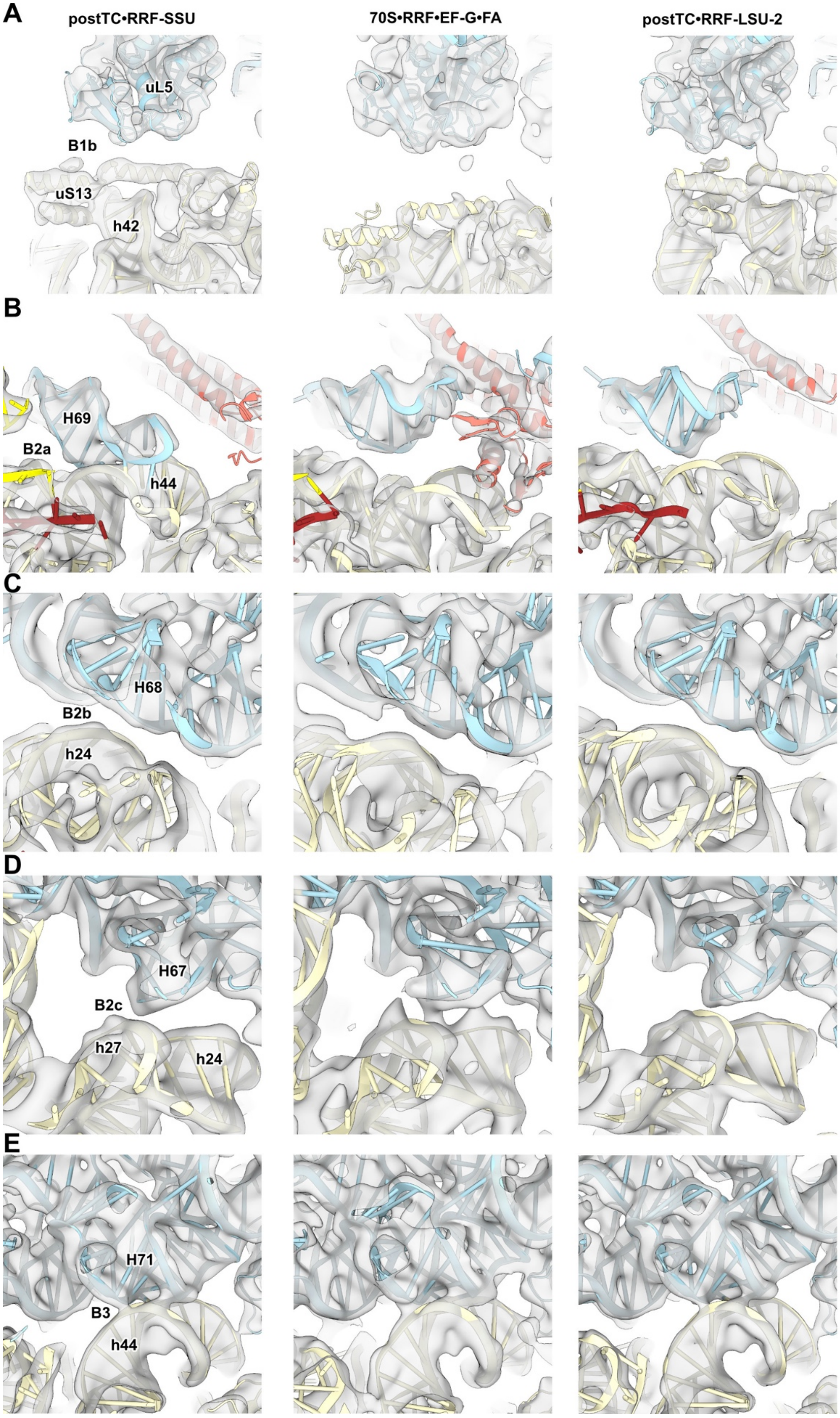
A
postTC•RRF-SSU
70S•RRF•EF-G•FA
postTC•RRF-LSU-2
uL5
B1b
uS13
h42
B
H69
B2a
h44
C
H68
B2b
h24
D
H67
B2c
h27
h24
E
H71
B3
h44

**Supplementary Figure 9: 6 Å low-pass filtered maps of inter-subunit bridges B1-B3.** LSU (light blue), SSU (light yellow), RRF (salmon), tRNA (yellow) and mRNA (brown) are shown. Bridges B1a and B1c are not shown due to weak density in postTC•RRF-SSU. **(A)** Bridge B1b. **(B)** Bridge B2a. **(C)** Bridge B2b. **(D)** Bridge B2c. **(E)** Bridge B3.

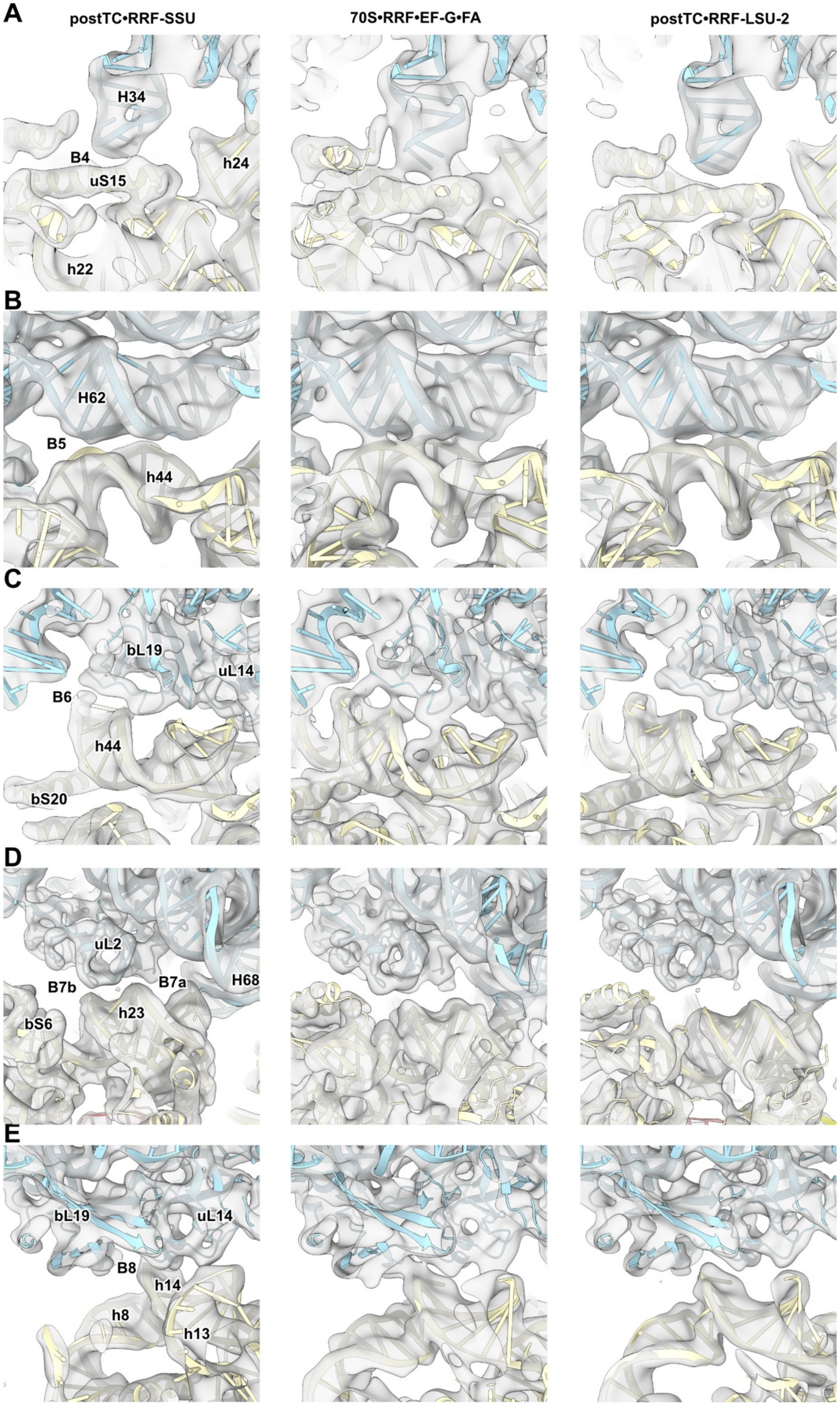
A
postTC•RRF-SSU
70S•RRF•EF-G•FA
postTC•RRF-LSU-2
H34
B4
uS15
h24
h22
B
H62
B5
h44
C
bL19
uL14
B6
h44
bS20
D
uL2
B7b
B7a
H68
h23
bS6
E
bL19
uL14
B8
h14
h8
h13

**Supplementary Figure 10: 6 Å low-pass filtered maps of inter-subunit bridges B4-B8.** LSU (light blue), SSU (light yellow), RRF (salmon), tRNA (yellow) and mRNA (brown) are displayed. **(A)** Bridge B4. **(B)** Bridge B5. **(C)** Bridge B6. **(D)** Bridges B7a and B7b. **(E)** Bridge B8.

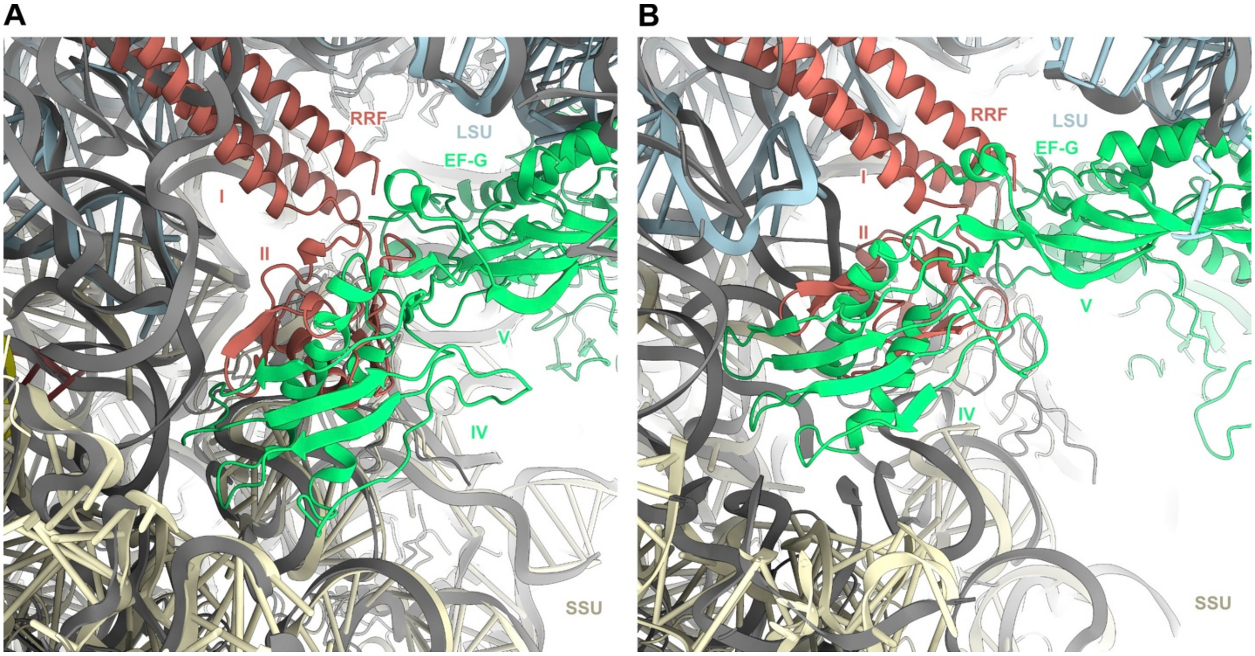


**Supplementary Figure 11: Steric clash between RRF and EF-G in different states of translocation. (A)** Superimposition by 23S rRNA of postTC•RRF-SSU (LSU, light blue; SSU, light yellow; RRF, salmon; tRNA, yellow; mRNA, brown) with the EF-G•GDP-Pi translocation intermediate (PDB 7SSL (4); ribosome and tRNA in gray; EF-G in green). Domain IV of EF-G in the EF-G•GDP-Pi state would sterically clash with domain II of RRF. **(B)** Superimposition by 23S rRNA of 70S•RRF•EF-G•FA (colors as (A), EF-G from this structure is not displayed) with the EF-G GDP-state on the ribosome (PDB 7SSD (4), ribosome and tRNA in gray, EF-G in green). Domain IV of EF-G•GDP would sterically clash with domain II of RRF and thus its conformation is different to 70S•RRF•EF-G•FA.
